# USP1 deubiquitinates protein kinase Akt to inhibit PI3K-Akt-FoxO signaling

**DOI:** 10.1101/654921

**Authors:** Dana Goldbraikh, Danielle Neufeld, Yara Mutlak-Eid, Inbal Lasry, Anna Parnis, Shenhav Cohen

## Abstract

PI3K-Akt-FoxO-mTOR signaling is the central pathway controlling growth and metabolism in all cells. Activation of this pathway requires ubiquitination of Akt prior to its activation by phosphorylation. Here, we found that the deubiquitinating (DUB) enzyme USP1 removes K63-linked polyubiquitin chains on Akt to sustain PI3K-Akt-FoxO signaling low during prolonged starvation. DUB screening platform identified USP1 as a direct DUB for Akt, and USP1 depletion in atrophying muscle increased Akt ubiquitination, PI3K-Akt-FoxO signaling, and glucose uptake during fasting. Co-immunoprecipitation and mass spectrometry identified Disabled-2 (Dab2) and the tuberous sclerosis complex TSC1/TSC2 as USP1 bound proteins. During starvation, Dab2 was essential for Akt recruitment to USP1/UAF1 complex, and for PI3K-Akt-FoxO inhibition. Additionally, to maintain its own protein levels high, USP1 limits TSC1 levels to sustain mTOR-mediated basal protein synthesis rates. This USP1-mediated suppression of PI3K-Akt-FoxO signaling probably contributes to insulin resistance in catabolic diseases and perhaps to malignancies seen with USP1 mutations.

## INTRODUCTION

Phosphoinositide 3-kinase (PI3K)–Akt–mammalian target of rapamycin (mTOR) signaling (Latres et al., 2005; Bonaldo and Sandri, 2013) is the central pathway controlling cell growth, proliferation, and metabolism. Activation of this pathway by IGF-I or insulin promotes cell division, and in non-dividing muscle cells, it promotes growth by stimulating overall protein synthesis and inhibiting protein degradation (Glass, 2005; Zhao et al., 2008; Mammucari et al., 2007). Conversely, inhibition of this pathway reduces cell survival and, in muscle, causes atrophy. A critical player at the core of PI3K-Akt signaling is the serine/threonine kinase Akt, which serves as an indispensable conduit for transmission of growth and survival signals from cell surface receptors. Because dysregulation of Akt results in various pathologies including muscle atrophy, cancer, insulin resistance (as occurs in type-2 diabetes or obesity), and neurological disorders, its activity must be tightly regulated in all cells.

Akt1 (referred to as Akt in this study) is a member of the Protein Kinase B (PKB) family of kinases, which comprises three isoforms in mammalian cells, Akt1/PKBα, Akt2/PKBβ, and Akt3/PKBγ. These isoforms are composed of an N-terminal pleckstrin homology (PH) domain, a central catalytic domain containing a T308 phosphorylation site, and a C-terminal regulatory domain containing a S473 phosphorylation site (Hanada et al., 2004). The kinases PDK-1 and mTOR2 phosphorylate Akt at T308 and S473, respectively (Boucher et al., 2014), downstream of PI3K. Phosphorylation of Akt at these residues is considered rate limiting and obligatory for maximal activation of Akt. Once activated, Akt inhibits the transcription factor FoxO by phosphorylation, preventing its nuclear translocation and stimulation of expression of atrophy-related genes (“atrogenes”)(Glass, 2005; Lecker et al., 2004). Akt also inhibits the negative regulator of mTORC1, the Tuberous Sclerosis Complex 1 and 2 (TSC1/2) (DeYoung et al., 2008), consequently leading to mTORC1-mediated phosphorylation of Ribosomal protein 6 Kinase β-1 (S6K1) (Magnuson et al., 2012; Thoreen et al., 2012) and activation of protein synthesis and cell growth (Amirouche et al., 2009). In addition, Akt activates glycogen synthesis by phosphorylating and inactivating Glycogen Synthase Kinase β (GSK3-β), which under low insulin conditions, inhibits glycogen synthesis (Eskelinen, 2006). When blood glucose and insulin levels rise, insulin promotes glucose uptake by enhancing trafficking of the glucose transporter GLUT4 to the plasma membrane (Ploug et al., 1998). However, during fasting and catabolic diseases, PI3K–Akt–mTOR signaling decreases, and consequently glucose uptake and protein synthesis fall; simultaneously, proteolysis increases largely through the FoxO-mediated expression of the atrogene program (Latres et al., 2005). In fact, activation of FoxO3 alone is sufficient to trigger proteolysis via the ubiquitin proteasome system (Sandri et al., 2004) and autophagy (Mammucari et al., 2007; Zhao et al., 2007), and to cause substantial atrophy (Sandri et al., 2004). Overproduction of Insulin-like Growth Factor 1 (IGF1) or Akt in mice, either through transgenic expression or by electroporation into muscles, was sufficient to reduce muscle weight loss, and induce systemic hypertrophy (Lai et al., 2004; Sacheck et al., 2004). These anabolic effects are mediated by Akt; however, the aberrant activation of this kinase and PI3K-Akt signaling may relay proliferative and pro-survival signals that are often associated with solid and hematological malignancies in human (Chen et al., 2011b; Burke et al., 2011). Because Akt controls diverse biological processes, from muscle growth and cell proliferation to survival and migration, its activity must be tightly regulated in normal cells. Previous studies in mouse embryonic fibroblasts proposed a new mode of regulation of Akt by ubiquitination, which appears to be required for Akt phosphorylation and activation (Yang et al., 2009). The present studies were undertaken to identify the enzyme that deubiquitinates Akt and suppresses PI3K-Akt signaling.

These studies have identified a novel regulator of Akt activity, the Ubiquitin-Specific Protease 1 (USP1). USP1 is a member of the USP family of cysteine proteases, which process ubiquitin or ubiquitin chains to reverse protein modification by ubiquitination. Mammalian cells contain over 50 members of this family, all of which have a catalytic core composed of conserved N-terminal Cys box motif and C-terminal His box motif (Hu et al., 2002). USP1 best characterized functions are in the nucleus as a regulator of DNA damage response, mainly in the Fanconi Anemia pathway (Nijman et al., 2005), and as a negative regulator of certain transcription factors to prevent cell differentiation (Williams et al., 2011). Recent evidence indicate an additional role for this enzyme in diabetes by promoting apoptosis of insulin secreting pancreatic β-cells (Maedler et al., 2018). Previously, a USP1 knockout mouse has been described that exhibits multiple developmental defects including, osteopenia (Williams et al., 2011), perinatal lethality, male infertility, chromosome instability, and a Fanconi Anemia phenotype (Kim et al., 2009). Although USP1 important roles in DNA repair mechanisms are well-documented, its precise functions in regulating metabolism and growth are not yet known. We demonstrate here that during fasting-induced atrophy, USP1 suppresses PI3K-Akt signaling through effects on the key component in this pathway, Akt.

Activation of USP1 requires association with the WD40 repeat containing protein USP1-associated factor 1 (UAF1) (Cohn et al., 2007), and its inhibition is mediated by phosphorylation (Cotto-Rios et al., 2011), by decreasing its protein levels via reduced gene expression (Williams et al., 2011), or by autocleavage at an internal diglycine motif (Gly670-Gly671) that promotes USP1 degradation by the proteasome (Piatkov et al., 2012). USP1 levels are elevated in several human cancers (Williams et al., 2011), where it seems to sustain DNA repair mechanisms; thus, inhibition of this enzyme may help hypersensitize cancer cells to chemotherapy-induced DNA damage (Mistry et al., 2013). Interestingly, mutations in USP1 have also been identified in certain human cancers (García-Santisteban et al., 2013), but the functional consequences of these mutations on USP1 activity remain elusive.

We demonstrate here a new role for USP1 in the inhibition of signaling by the PI3K-Akt cascade. USP1 directly deubiquitinates Akt leading to suppression of Akt phosphorylation and inactivation. Recruitment of Akt to USP1/UAF1 complex is mediated by the tumor suppressor Disabled-2 (Dab2), which harbors Akt binding domain, and whose ectopic expression in cancer cell lines is sufficient to inhibit PI3K-Akt signaling and cell proliferation (Yang et al., 2019; Koral and Erkan, 2012; Zhoul et al., 2005). In order to maintain its own protein levels high, USP1 also limits the protein content of TSC1 to sustain basal rates of protein synthesis. Thus, USP1 functions as a novel inhibitor of Akt, which is important for regulation of PI3K-Akt signaling and maintenance of cellular homeostasis and normal muscle size.

## RESULTS

### USP1 is a deubiquitinating enzyme for Akt

Because PI3K-Akt-FoxO-mTOR pathway is the primary regulator of cell growth and metabolism, and given the importance of Akt ubiquitination for its activity (Yang et al., 2009), we set out to identify the specific DUB that reduces Akt ubiquitination and consequently inhibits PI3K-Akt signaling. To initially confirm that Akt is actually deubiquitinated when PI3K-Akt-FoxO-mTOR falls, we studied mouse muscles under the physiological condition of fasting, when blood glucose and insulin levels are low and PI3K-Akt-FoxO-mTOR signaling in all cells is inhibited. Analysis of skeletal muscle extracts from fed and fasted (2d) mice by SDS-PAGE and immunoblotting showed that Akt is ubiquitinated under normal conditions but its ubiquitination levels are reduced during fasting, while the unmodified form of Akt accumulates (Fig. 1A). These findings suggested that Akt is deubiquitinated *in vivo* during fasting.

**Figure 1.**
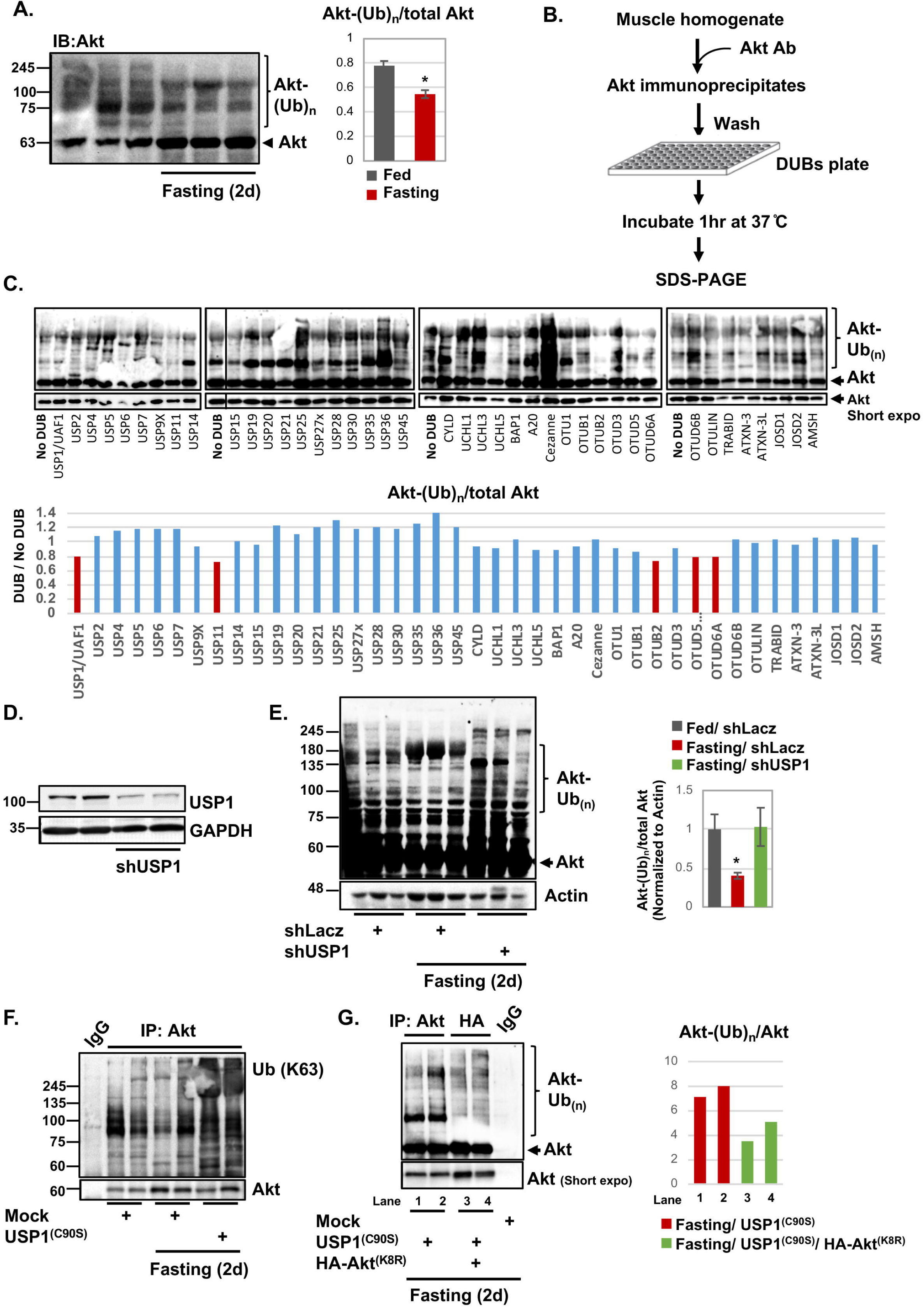
USP1 is a deubiquitinating enzyme for Akt. (A) Akt is deubiquitinated during fasting. Left: Soluble fractions of TA muscles from fed and fasted mice were analyzed by SDS-PAGE and immunoblot using Akt antibody. Right: densitometric measurement of presented blots (n=3). * p<0.05 vs. fed. (B) A scheme for DUBs screening experiment to identify the enzyme that directly deubiquitinates Akt *in vitro*. (C) Ubiquitinated Akt is a substrate for USP1 *in vitro*. Top: Akt was immunoprecipitated from the soluble fraction of TA muscle from fed mice, and was subjected to an *in vitro* deubiquitination by a panel of purified DUBs arrayed in a multi-well plate. The dividing line indicates the removal of an intervening lane for presentation purposes. Bottom: Densitometric measurements of presented blots. Data is presented as the ratio between ubiquitinated Akt to total Akt in each well. (D) shRNA-mediated knockdown of USP1 in Hela cells. Soluble extracts were analyzed by immunoblotting. (E) USP1 deubiquitinates Akt during fasting *in vivo*. Left: Soluble fractions of TA muscles transfected with shLacz control or shUSP1 from fed and fasted mice were analyzed by SDS-PAGE and immunoblotting using anti-Akt. The actin blot serves as a loading control. Right: densitometric measurements of presented blots (n=3). * p<0.05 vs. fed. (F) USP1 removes K63-linked polyubiquitin chains on Akt *in vivo*. Akt was immunoprecipitated from the soluble fraction of muscles expressing USP1^(C90S)^ or control plasmids. Mouse IgG was used as a control for non-specific binding. Protein precipitates were subjected to immunoblotting with an antibody against K63-linked polyubiquitin conjugates. (G) USP1 removes polyubiquitin chains linked to K8 on Akt. Akt was immunoprecipitated from the soluble fraction of TA muscles transfected with USP1^(C90S)^, HA-Akt^(K8R)^ or control from fasted mice, and protein precipitates were analyzed by immunoblotting with anti-Akt antibody. Mouse IgG was used as a control for non-specific binding. Right: Densitometric measurements of presented blots. Data is presented as the ratio between ubiquitinated Akt to total Akt in each lane (n=2).

To identify the DUB that can directly deubiquitinate Akt *in vitro*, we immunoprecipitated Akt from normal muscle homogenate, and added the precipitates to a DUB-screening plate containing an array of pure active DUBs (Fig. 1B). Five DUBs, USP1/UAF1, USP11, OTUB2, OTUD5 (p177S), and OTUD6A, cleaved the polyubiquitin chain on Akt and reduced its ubiquitination levels by at least 20% compared with ubiquitinated Akt in reactions that did not contain a DUB (Fig. 1C). Among these DUBs, USP1 was deemed the most intriguing candidate as the enzyme deubiquitinating and inhibiting Akt because it preferentially removes K63-linked chains (Chen et al., 2011a), the type of polyubiquitination that is essential for Akt activation (Yang et al., 2009), and it is the only DUB among the five that has been linked to diabetes, a pathological condition associated with dysregulated insulin-PI3K-Akt signaling (Maedler et al., 2018).

To determine whether USP1 is a deubiquitinating enzyme for Akt *in vivo*, we downregulated this enzyme in mouse muscle by the electroporation of a specific shRNA (shUSP1), which can efficiently reduce USP1 content (Fig. 1D), and analyzed the effects on Akt ubiquitination levels during fasting. As shown above (Fig. 1A), analysis of the soluble fraction from muscles expressing control plasmid (shLacz) from fed and fasted mice indicated that upon fasting there was a decrease in high molecular weight ubiquitinated species of Akt (fig. 1E). However, downregulation of USP1 blocked this decrease in Akt ubiquitination levels, and instead, Akt accumulated as ubiquitinated species (Fig. 1E). Similar results were obtained when we inhibited USP1 by a different approach, i.e. by the electroporation of a GFP-tagged USP1 dominant negative encoding plasmid (USP1^(C90S)^)(Olazabal-Herrero et al., 2016), which lacks the catalytic cysteine and thus can bind substrates but cannot deubiquitinate them (Fig. 1F). Immunoprecipitation of Akt from USP1^(C90S)^ expressing muscles from fasted mice, and analysis of protein precipitates by SDS-PAGE and immunoblotting using anti K63-linked ubiquitin chains antibody revealed that in the muscles lacking functional USP1 (expressing USP1^(C90S)^) Akt accumulated as K63 ubiquitinated protein (Fig. 1F). In fact, the content of K63 ubiquitinated Akt in these muscles exceeded the amounts observed in muscles of fed mice (P < 0.05), even though only about half the muscle fibers were transfected, suggesting that K63 polyubiquitination of Akt may increase during fasting, and USP1 catalyzes the deubiquitination of this protein.

Akt ubiquitination on K8 within the PH domain is essential for its activation (Yang et al., 2009). To learn whether USP1 cleaves the ubiquitin chain that is linked to K8 on Akt, we co-electroporated muscles with USP1^(C90S)^ to cause accumulation of ubiquitinated Akt, and either shLacz or a plasmid encoding HA-tagged Akt carrying a K8-to-R mutation (HA-Akt^(K8R)^)(Fig. 1G). By 2d of fasting, HA-Akt^(K8R)^ immunoprecipitated from transfected muscles showed limited ubiquitination, i.e., much less than the endogenous Akt immunoprecipitated from muscles expressing USP1^(C90S)^ alone (Fig. 1G, compare lanes 1-2 with 3-4). These findings indicate that on inhibition of USP1 during fasting, Akt is ubiquitinated on K8. Thus, USP1 is essential for Akt deubiquitination *in vivo*, which most likely leads to inhibition of PI3K-Akt signaling.

### Deubiquitination by USP1 reduces Akt phosphorylation and PI3K-Akt-FoxO signaling

To determine whether USP1-mediated Akt deubiquitination in fact influences PI3K–Akt– FoxO signaling, we inhibited USP1 by electroporation of USP1^(C90S)^ into mouse muscle during fasting. By two days of fasting, phosphorylation of insulin receptor, Akt, and its targets FoxO3 and GSK3β, was markedly reduced (Fig. 2A) (Sandri et al., 2004; Stitt et al., 2004). However, inhibition of USP1 almost completely blocked this response to fasting. In fact, the levels of phosphorylated Akt (T308), FoxO3, and GSK3β were similar to those in muscles from fed mice (Fig. 2A). Normally during fasting, FoxO is activated (dephosphorylated) and stimulates the expression of atrogenes, including the ubiquitin ligases MuRF1 and Atrogin1, which are essential for rapid fiber atrophy (Fig. 2B)(Bodine et al., 2001; Gomes et al., 2001). However, USP1 downregulation with shRNA (shUSP1) resulted in a marked decrease in MuRF1 and Atrogin1 expression in the TA muscles during fasting (Fig. 2B). This inhibition of atrogene expression during fasting together with the maintenance of normal PI3K–Akt–mTOR signaling should block muscle wasting (Cohen et al., 2014; Aweida et al., 2018). Accordingly, the mean cross-sectional area of 500 fibers expressing shUSP1 (and GFP to identify transfected fibers) was bigger than that of 500 non-transfected fibers (Fig. 2C). Moreover, by 2 d of fasting, there was a 31% decrease in the mean weight of the control TA muscles below levels in fed mice (Fig. 2D), but USP1 downregulation clearly attenuated this wasting, even though ∼30% of the fibers were not transfected with the shUSP1 (Fig. 2D). In addition to regulating cell size and protein balance, the PI3K–Akt–FoxO pathway also mediates insulin’s stimulation of glucose uptake into muscle and adipose tissue. As predicted, inhibition of USP1 by the injection of mice with the USP1 specific inhibitor ML323 (Liang et al., 2014), significantly improved glucose tolerance during fasting (Fig. 2E). Thus, USP1 function is critical in causing the reduction in PI3K–Akt–FoxO signaling during fasting, that triggers the decrease in glucose uptake and protein synthesis, the FoxO-mediated expression of the atrogene program, and muscle wasting (Cohen et al., 2014; Aweida et al., 2018).

**Figure 2.**
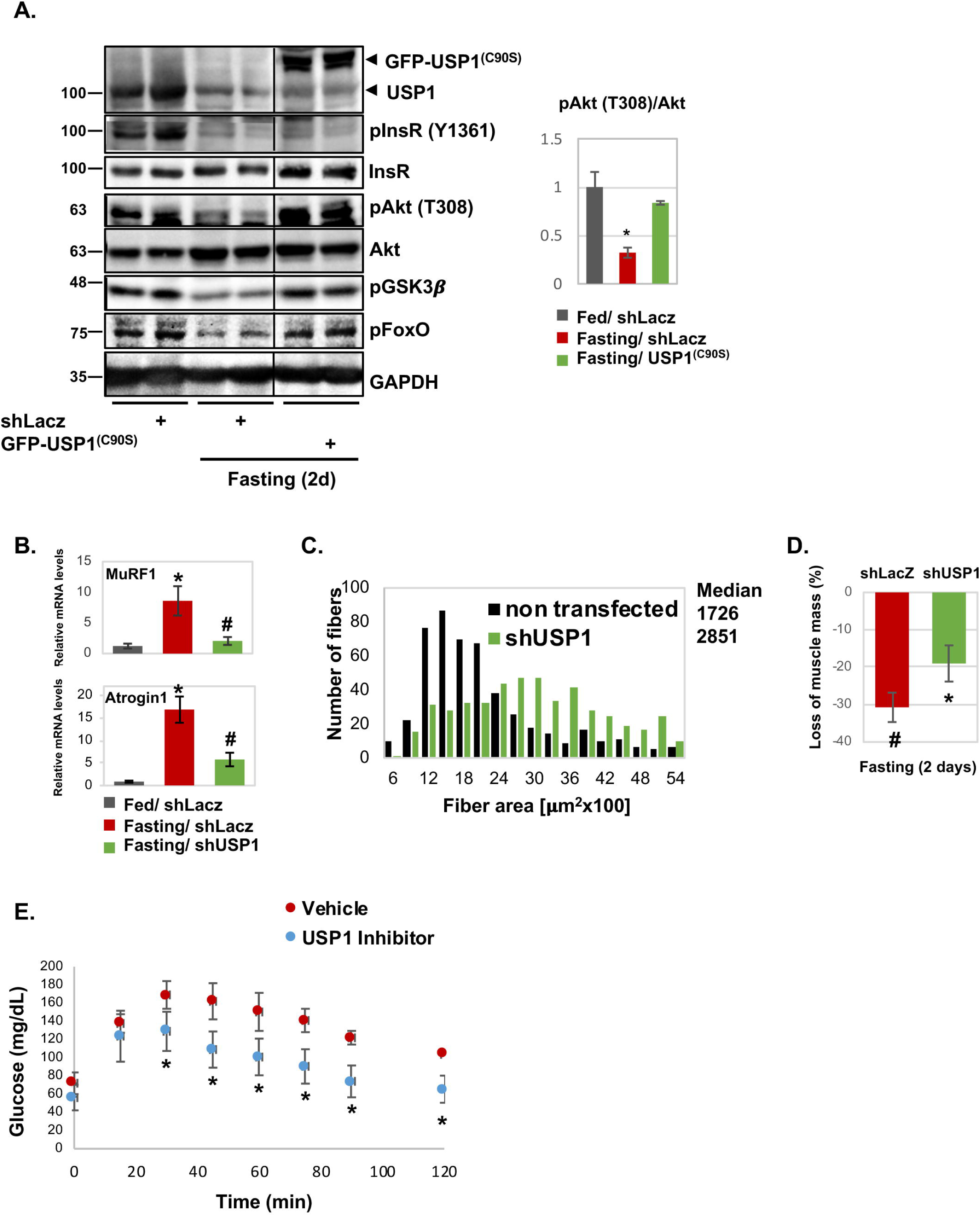
Deubiquitination by USP1 reduces Akt phosphorylation and PI3K-Akt-FoxO signaling. (A) Inhibition of USP1 increases PI3K–Akt–FoxO signaling during fasting. Soluble fractions of normal and atrophying muscles expressing shLacz or USP1^(C90S)^ were analyzed by SDS-PAGE and immunoblot. The black line indicate the removal of intervening lanes for presentation purposes. Right: densitometric measurements of presented pAkt (T308) and Akt blots. Data is presented as ratio of phosphorylated Akt to total Akt (n=3). * p<0.05 vs. fed. (B) Downregulation of USP1 reduces MuRF1 and Atrogin1 expression during fasting. Quantitative RT-PCR of mRNA preparations from atrophying and control muscles expressing shLacz or shUSP1 using primers for MuRF1 and Atrogin1. Data are plotted as the mean fold change relative to fed control. n = 4. *, P < 0.05 vs. shLacz in fed. #, P < 0.05 vs. shLacz in fasting. (C) Downregulation of USP1 markedly reduces muscle fiber atrophy. Cross-sectional areas of 500 fibers transfected with shUSP1 (that express GFP, green bars) vs. 500 nontransfected fibers (black bars) in the same muscle. n = 5 mice. (D) Downregulation of USP1attenuates the loss of muscle mass during fasting. shUSP1 was delivered to more than 60% of muscle fibers. Mean weights of electroporated muscles are plotted as the percent weight loss. n = 6. #, P < 0.05 vs. shLacz in fed; *, P < 0.05 vs. shLacz in fasting. (E) USP1 inhibition significantly increases glucose tolerance in mice during fasting. Mice injected i.p. with specific USP1 Inhibitor (12ug/gr body weight) or saline during fasting (2d) were subjected to glucose tolerance test. Blood glucose levels were measured at the indicated time points following glucose injection (1 mg/gr body weight). Data is depicted in a graph as mg/dL glucose (n=3). * p<0.05 vs. mice injected with saline.

### Akt is recruited to USP1/UAF1 complex in fasting

To identify the growth regulatory factor that recruits Akt to USP1, we performed in parallel immunoprecipitation of components that interact with Akt, i.e. USP1 and its cofactor UAF1, and FoxO3, and identified the bound proteins by mass spectrometry. Muscle homogenates from fasted mice, where Akt is dephosphorylated and inactivated, were incubated with specific antibodies against USP1, UAF1 or FoxO3, or with a non-specific IgG, and protein precipitates were washed extensively with buffer containing 500 mM NaCl to remove nonspecific or weakly associated proteins (Fig. 3A). Mass spectrometry analysis identified about 80 proteins that separately bound USP1, UAF1, or FoxO3, and were at least three-fold more abundant in the immunoprecipitation samples than in IgG control (Table S1). Among these proteins, only 43 were common binding partners of all three components (Fig. 3B and Table S1), and among them only 29 proteins were at least 1000 times more abundant in the immunoprecipitation samples than in IgG control. These common proteins included TSC1/TSC2 complex, which lies downstream of PI3K-Akt signaling (Wullschleger et al., 2006; Inoki et al., 2002), and Dab2 that can bind Akt and regulate PI3K-Akt signaling in epithelia (Yang et al., 2019; Koral and Erkan, 2012). Akt was not identified by mass spectrometry even though it is clearly a substrate for USP1 (Figs. 1-2).

**Figure 3.**
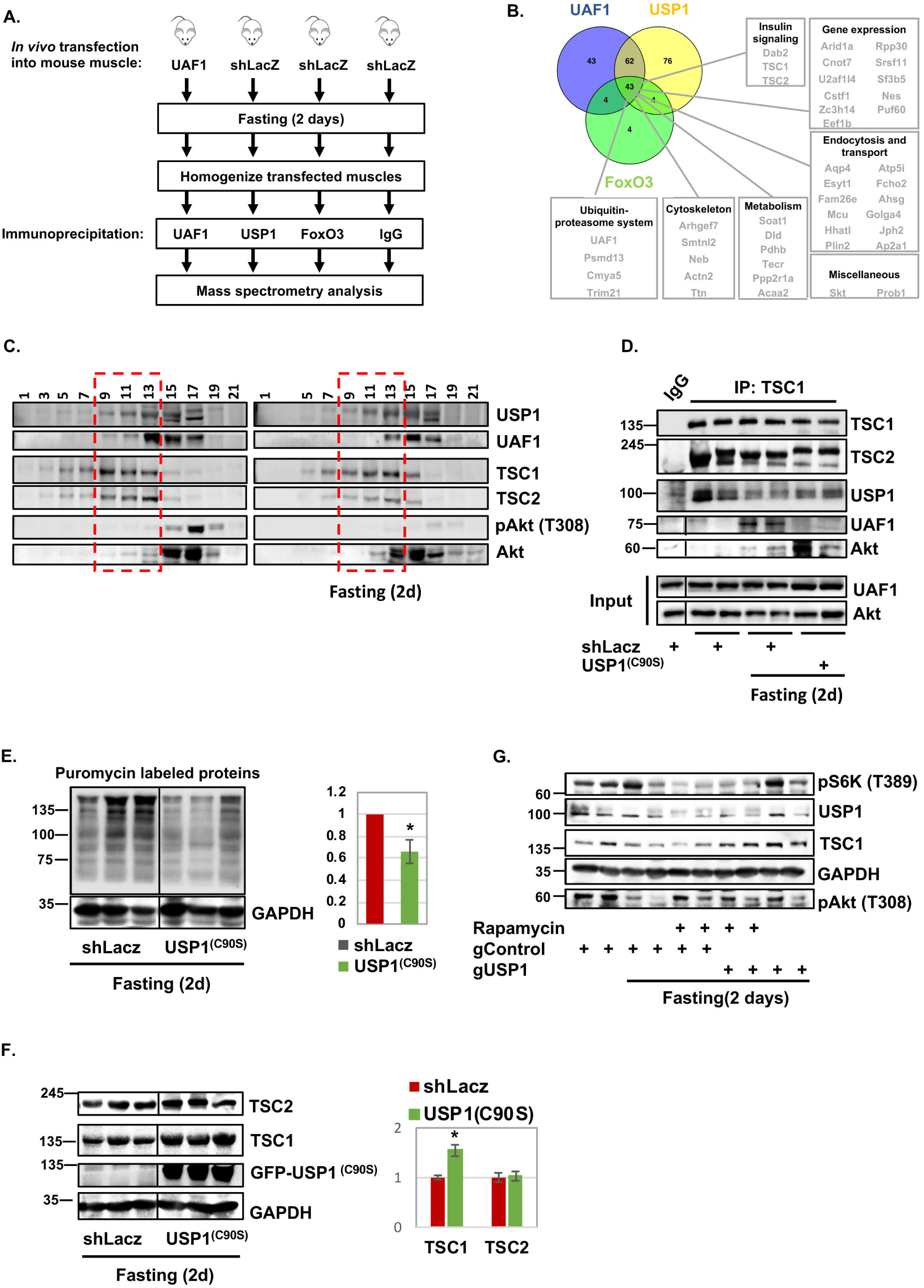
Akt is recruited to USP1-TSC1 complex in fasting. (A) Scheme of affinity purification technique to isolate components that bind Akt. Three immunoprecipitation experiments of USP1, UAF1 and FoxO from muscle homogenates during fasting were performed in parallel to non-specific mouse IgG control, and protein precipitates were analyzed by mass spectrometry. (B) Mass spectrometry data obtained in (A) is presented using Venny diagram tool. Only proteins that were at least threefold more abundant in the immunoprecipitation samples than in IgG control were considered. (C) In fasting, Akt sediments to glycerol gradient fractions containing USP1/UAF1 and TSC1/TSC2. Muscle homogenates from fed or fasted mice were separated on 10%-40% glycerol gradient, and alternate fractions were analyzed by SDS-PAGE and immunoblot. (D) During fasting, Akt is recruited to USP1-TSC1 complex. TSC1 was immunoprecipitated from the soluble fraction of muscles expressing shLacz or USP1^(C90S)^ from fed or fasted mice. Mouse IgG was used as a control for non-specific binding. Precipitates were analyzed by immunoblotting. Black lines indicate the removal of intervening lanes for presentation purposes. (E) During fasting, inhibition of USP1 reduces rates of protein synthesis. Mice were injected with puromycin, and soluble fractions of electroporated muscles were analyzed by immunoblotting using puromycin antibody. Right: Densitometric measurement of the presented blot. n = 3. *, P < 0.05 vs. shLacz. (F) Inhibition of USP1 results in TSC1 accumulation. Soluble fraction of muscles expressing shLacz or USP1^(C90S)^ from fasted mice were analyzed by SDS-PAGE and immunoblotting. Right: Densitometric measurement of the presented blot. n = 3. *, P < 0.05 vs. shLacz. Data is presented as protein content (AU) in USP1^(C90S)^ vs. shLacz. (G) mTOR inhibition with rapamycin results in reduced USP1 protein content and enhanced Akt phosphorylation at T308. However, a simultaneous downregulation of USP1 is required to promote TSC1 accumulation. Soluble fraction of muscles expressing shLacz or USP1^(C90S)^ from fed and fasted mice injected i.p. with rapamycin (6mg/kg body weight) or saline were analyzed by SDS-PAGE and immunoblot.

Because TSC1/TSC2 complex is a direct substrate of Akt (Inoki et al., 2002), we initially determined if these proteins influence Akt activity during fasting, and the role of USP1. Glycerol gradient fractionation of muscle homogenates from fed mice indicated that TSC1, TSC2, and USP1/UAF1 sedimented to the same fractions, as determined by the distribution of these proteins across the gradient (Fig. 3C). Interestingly, Akt was virtually absent in these fractions, and mainly sedimented to lighter fractions containing UAF1 and truncated mostly likely inactive USP1 (Huang et al., 2006)(Fig. 3C). In these lighter fractions, where USP1 is cleaved and is probably inactive, Akt was present in its phosphorylated active form (T308), which is consistent with USP1 playing an important role in reducing Akt phosphorylation (Fig. 2A). During fasting, however, Akt moved to the heavier fractions containing USP1/UAF1, and TSC1/TSC2, and its phosphorylation was reduced (Fig. 3C).

To determine if these proteins physically interact *in vivo* and whether Akt is in fact recruited to this protein assembly in fasting, we performed immunoprecipitation experiments from muscles expressing GFP-USP1^(C90S)^ or control, from fed or fasted mice. TSC2 and USP1 could be coprecipitated with TSC1 from muscles of fed or fasted mice (Fig. 3D). However, association of UAF1 and Akt with this protein assembly was evident only during fasting (Fig. 3D). It is noteworthy that neither USP1 nor UAF1 mRNA increased during fasting (Fig. S1), although USP1 is clearly essential for the reduction in Akt phosphorylation and PI3K-Akt signaling (Fig. 2). Its function probably increases by the enhanced association with UAF1 (Fig. 3D)(Cohn et al., 2007). Interestingly, USP1/UAF1 association does not seem to be required for Akt recruitment to USP1 because Akt remained bound to this protein even in muscles expressing USP1^(C90S)^ where USP1/UAF1 association was perturbed (Fig. 3D). Thus, Akt recruitment to USP1 probably requires an additional signal beyond USP1/UAF1 association. Finally, UAF1 failed to bind USP1-TSC1 complex in muscles overexpressing USP1^(C90S)^ (Fig. 3D) probably because the catalytically dead USP1 enzyme captures this protein away from the native functional complex.

Further studies determined whether USP1-TSC1 association shown above is important for Akt inactivation in fasting. Perhaps, USP1 by deubiquitinating and inhibiting Akt, promotes activation of TSC1/TSC2 and consequently reduces rates of protein synthesis. Alternatively, during fasting, USP1 may maintain its own protein levels high by limiting TSC1 accumulation (not through effects on Akt) to sustain basal rates of mTOR-mediated protein synthesis. To test these ideas, we initially inhibited USP1 in mouse muscle and analyzed the effects on rates of protein synthesis. After electroporation of USP1^(C90S)^ into TA muscle to inhibit USP1 during fasting, puromycin incorporation into newly translated proteins was lower than in shLacz-expressing muscles (Fig. 3E), and TSC1 protein levels were increased (Fig. 3F). This accumulation did not result from increased gene expression because the mRNA levels of TSC1 were similar in atrophying muscles expressing shLacz or USP1^(C90S)^ (Fig. S2A), suggesting that during fasting USP1 limits TSC1 content by promoting its degradation. Thus, USP1 sustains basal rates of protein synthesis by limiting TSC1 accumulation in fasting. Accordingly, inhibition of mTOR in fasting by the injection of mice with rapamycin, as shown by lower amounts of phosphorylated S6K (mTOR-downstream target)(Fig. 3G), caused a decrease in USP1 protein levels, which can account for the dramatic increase in phosphorylated Akt in these muscles (Fig. 3G). Moreover, downregulation of USP1 by the co-electroporation of Cas9 encoding plasmid and USP1 guiding RNAs resulted in increased Akt phosphorylation and caused accumulation of TSC1, with or without a simultaneous treatment with rapamycin (Fig. 3G). Similar results were obtained by the downregulation of USP1 with shUSP1 (Fig. S2B). Possibly, a reduction in USP1 protein levels below a critical threshold is necessary to cause TSC1 accumulation in fasting. Thus, when PI3K-Akt signaling is low, USP1 limits TSC1 protein levels to promote basal rates of mTOR-mediated protein synthesis and sustain its own protein levels high. A similar accumulation of TSC1 was observed by the downregulation of USP1 after 24hr and 36hr of food deprivation, even before any effects on Akt phosphorylation could be observed (Fig. S2C), suggesting that USP1 function toward TSC1 is independent from its role in reducing Akt activity.

### Dab2 facilitates Akt recruitment to USP1-TSC1 complex

Together, these observations suggested that Akt is recruited to USP1-TSC1 complex by an additional factor. In cancer cells, overexpression of Dab2 reduces PI3K-Akt signaling, and Dab2 C-terminal proline rich domain can bind Akt (Yang et al., 2019; Koral and Erkan, 2012). Because Dab2 in muscle extracts was bound to USP1/UAF1, and other Akt interacting partners including FoxO (Fig. 3B and Table S1), we initially determined whether it influences PI3K-Akt activity during fasting. For this purpose, we used a vector encoding the C-terminal proline rich domain of Dab2 (Dab2-DN), which functions as a dominant-negative inhibitor of Dab2-Akt association (Koral and Erkan, 2012). Because this domain in Dab2 is normally recognized by Akt (Koral and Erkan, 2012), the expression of the truncated mutant should compete with the endogenous Dab2 on association with Akt. We electroporated TA muscles from fed or fasted mice with control plasmid or the Dab2-DN, and analyzed the transfected muscles by immunoprecipitation. As shown above (Fig. 3D), Akt association with USP1 and TSC1 was greater during fasting than in muscles of fed mice (Fig. 4A). Similarly, Dab2 association with this protein assembly increased during fasting (Fig. 4A), when Akt is deubiquitinated (Fig. 1) and PI3K-Akt signaling falls (Fig. 2A). This change in Akt association in fasting seemed to require Dab2 because inhibition of Dab2-Akt association by the expression of Dab2-DN blocked this recruitment of Akt and Dab2 to USP1-containing complex (Fig. 4A). By competing with the endogenous Dab2 on association with Akt, this truncated mutant seems to also capture the endogenous Dab2 and prevent its association with the functional USP1-TSC1 complex (Fig. 4A). Thus, during fasting, Dab2 enhances the association between Akt and USP1.

**Figure 4.**
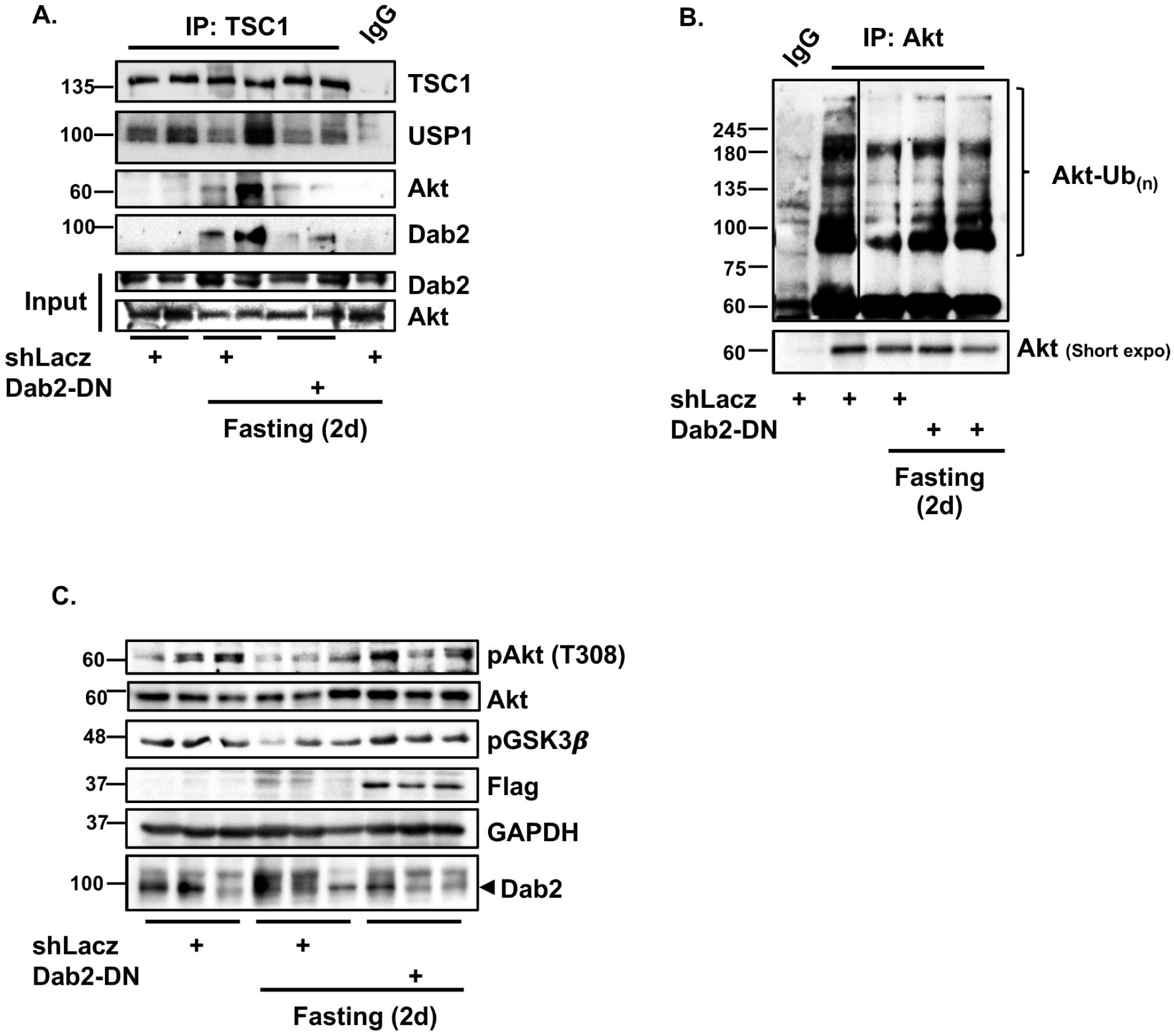
Dab2 facilitates Akt recruitment to USP1-TSC1 complex. (A) During fasting, inhibition of Dab2-Akt association reduces the interaction of Akt with USP1-TSC1 complex. TSC1 was immunoprecipitated from the soluble fraction of muscles expressing shLacz or Dab2-DN from fed or fasted mice. Mouse IgG was used as a control for non-specific binding. Precipitates were analyzed by immunoblotting. (B) Dab2 promotes Akt deubiquitination *in vivo*. Akt was immunoprecipitated from the soluble fraction of muscles expressing shLacz or Dab2-DN from fed and fasted mice using a specific antibody. Mouse IgG was used as a control for non-specific binding. Protein precipitates were analyzed by Western blot analysis using Akt antibody. (C) Inhibition of Dab2-Akt association enhances Akt phosphorylation in fasting. Soluble fraction of TA muscles transfected with shLacz or Dab2-DN from fed and fasted mice were analyzed by SDS-PAGE and immunoblotting using the indicated antibodies.

Because Dab2 mediates Akt recruitment to USP1 during fasting, reducing its association with Akt by the expression of Dab2-DN should prevent Akt deubiquitination and activate PI3K–Akt–FoxO signaling. To determine if Dab2 facilitates Akt deubiquitination, we immunoprecipitated Akt from muscles from fed and fasted mice, and analyzed by SDS-PAGE and immunoblotting using Akt antibody. As shown above, the high molecular weight ubiquitinated species of Akt were reduced in fasting (Fig. 4B), where Dab2 is functional and facilitates the recruitment of Akt to USP1 (Fig. 4A). This deubiquitination of Akt is dependent on Dab2 because reducing Dab2 function by the expression of Dab2-DN resulted in accumulation of high molecular weight conjugates of Akt (Fig. 4B). In fact, this accumulation of ubiquitinated Akt by Dab2-DN must underestimate the protective effects of Dab2 inhibition because only about half the fibers were transfected. In addition, Akt phosphorylation decreased by two days fasting, when its association with USP1 increased (Fig. 3D); however, this effect was markedly attenuated in muscles expressing Dab2-DN (Fig. 4C) because in these muscles Akt recruitment to USP1 was reduced (Fig. 4A). It is noteworthy that neither Dab2 protein nor mRNA increased upon fasting (Figs. S3 and 4C), although this protein is clearly essential for Akt deubiquitination and for suppression of PI3K–Akt–FoxO signaling when blood insulin levels are low (Fig. 4A-C). Thus, Dab2 appears critical for the USP1-induced reduction in Akt ubiquitination in fasting, and for the resulting inhibition of Akt and PI3K-Akt-FoxO signaling.

## DISCUSSION

We uncovered a novel mechanism that suppresses PI3K-Akt signaling, and involves the deubiquitination of Akt by USP1. Because USP1 inhibition during fasting increased PI3K– Akt–FoxO signaling, enhanced glucose uptake, and blocked muscle atrophy (Fig. 2), USP1-mediated inhibition of Akt function may also contribute to insulin resistance and muscle wasting in various catabolic states (e.g., diabetes, metabolic syndrome, or sepsis), and perhaps to the malignancies seen with USP1 mutations. Recent evidence indicate a role for this enzyme in diabetes by promoting apoptosis of insulin secreting pancreatic β-cells (Maedler et al., 2018), and we show here that USP1 also reduces tissue sensitivity to insulin. Consequently, USP1 may represent a new therapeutic target to block muscle wasting and the insulin resistance characteristic of many disease states (e.g., cancer cachexia, sepsis, and renal failure), and stimulating USP1 actions toward Akt in order to inhibit PI3K-Akt -FoxO signaling may have therapeutic benefits in the treatment of cancer.

Under normal conditions, Akt is ubiquitinated by K63-linked polyubiquitin chains, which is required for its activation and for PI3K-Akt signaling (Figs. 1-2)(Yang et al., 2009). In MEF cells, this ubiquitination is catalyzed by the ubiquitin ligase TRAF6, and appears to be required for Akt phosphorylation (Yang et al., 2009). TRAF6 also plays a role in muscle atrophy (Kumar et al., 2012), and may thus be involved in Akt ubiquitination in muscles depleted in active USP1 (Fig. 1E-G). As shown here, during fasting, cleavage of these K63 ubiquitin chains by USP1 caused loss of Akt phosphorylation at T308. By deubiquitinating Akt, USP1 may prevent Akt phosphorylation by PDK1, or may enhance Akt dephosphorylation at this site by certain phosphatases.

The systemic atrophy of muscle upon fasting is signaled by the low levels of insulin and IGF-1 and increased levels of glucocorticoids, which cause inhibition of PI3K-Akt signaling and accelerated proteolysis via FoxO-mediated expression of the atrogene program (Sandri et al., 2004). We show here, that in addition to the fall in insulin, PI3K-Akt activity is suppressed in prolonged starvation by USP1 to efficiently terminate the transmission of growth signals when cellular energy charge is low. Accordingly, loss of USP1 function by itself is sufficient to enhance PI3K-Akt activity, which prevents atrogenes expression, protein degradation and atrophy (Fig. 2). Under these conditions, protein synthesis rates are low due to the inhibition of mTOR by TSC1/TSC2 complex (Sandri, 2008). However, we surprisingly show that TSC1 protein levels are in fact restricted in fasting by USP1 in order to maintain basal rates of mTOR-mediated protein synthesis and sustain USP1 protein levels high enough to exert its effects. Accordingly, downregulation of USP1 resulted in accumulation of TSC1 in muscles from fasted mice (Fig. 3F), and reduced rates of protein synthesis (Fig. 3E). In addition, treatment of mice with the mTOR inhibitor, rapamycin, reduced USP1 protein levels and consequently Akt phosphorylation increased (Fig. 3G). Therefore, USP1-mediated loss of Akt phosphorylation does not involve the function of mTOR and occurs upstream, i.e. by the direct action of USP1 on Akt (Fig. 1C). Interestingly, in addition to rapamycin treatment, a simultaneous downregulation of USP1 by an *in vivo* gene manipulation was required to cause TSC1 accumulation, probably because the low USP1 protein levels in mice treated with rapamycin alone were sufficient to limit TSC1 protein levels. These effects appear to involve USP1-mediated regulation of TSC1 stability because TSC1 expression did not increase in muscle lacking functional USP1 (Fig. S3). In any case, maintenance of basal rates of protein synthesis in times of scarcity is crucial for survival, because the induction of enzymes that promote proteolysis is required to facilitate muscle protein degradation, and the amino acids produced are converted to glucose in the liver to nurture the brain.

We previously identified the ubiquitin ligase Trim32 as a novel negative regulator of PI3K-Akt signaling in fasting (Cohen et al., 2014), and the present studies demonstrate a critical role for the deubiquitinating enzyme USP1 in the inhibition of this pathway. Whether these proteins function together in regulating PI3K-Akt activity is an important question for future research. Trim32 influences PI3K-Akt activity by perturbing PI3K association with the desmosomal component plakoglobin (Cohen et al., 2014), while USP1 acts directly on Akt to inhibit its activity by deubiquitination. Because USP1 is not induced during fasting (Fig. S1), its activity most likely increases on binding to UAF1 (Fig. 3D). This heterodimerization represents the most critical event in activation of USP1 (Cohn et al., 2007), and appears to require USP1 catalytic activity (Fig. 3D). Interestingly, these proteins together with TSC1/TSC2 form one intact complex probably to efficiently exert the inhibitory effects on Akt to maintain PI3K-Akt signaling low. Surprisingly, however, Akt remained bound to TSC1/TSC2 and USP1/UAF1 even when UAF1 failed to bind USP1 in muscles lacking functional USP1 (i.e. overexpressing USP1 catalytically dead mutant)(Fig. 3D). These findings indicate that Akt association with USP1 is not dependent on USP1 activity, and the interaction between Akt and this enzyme is most likely mediated by additional associated proteins. In fact, we identified Dab2 as essential for Akt recruitment to USP1 in fasting. Dab2 can bind Akt also in kidney cells, where it regulates albumin endocytosis (Koral and Erkan, 2012). Moreover, in epithelia, Dab2 regulates signaling pathways (e.g. by Wnt, TGF-β) that control differentiation, cell proliferation, and motility (Yang et al., 2019; Zhoul et al., 2005; Hocevar et al., 2001; Jiang et al., 2012; Hocevar et al., 2005), and as shown here, it binds Akt in skeletal muscle and reduces signaling through the PI3K-Akt cascade during fasting. Activation of this pathway by IGF-I or insulin promotes muscle growth by enhancing overall protein synthesis and inhibiting protein degradation (Sacheck et al., 2004; Glass, 2005), and impaired signaling through this pathway in untreated diabetes, sepsis, and cancer cachexia (Zhou et al., 2010) can cause severe muscle wasting, which can be inhibited by the activation of PI3K–Akt–FoxO signaling (Wang et al., 2006). Conversely, inhibition of these kinases has a major therapeutic benefit in treating cancer by blocking cell proliferation. Interestingly, Dab2 has been shown to be an important regulator of cell proliferation because its loss causes cancer (Sheng et al., 2000; Yang et al., 2019), and its ectopic expression blocks cancer cells proliferation (Zhoul et al., 2005). As shown here, these growth suppressive effects by Dab2 activity are most probably due to enhanced Akt association with USP1 and the resulting inhibition of PI3K-Akt signaling. Clearly, the exact mode of regulation of this association and the specific domains in each protein that constitute these interactions merit further study. In fasting, Dab2-Akt association is mediated by Dab2’s C-terminal proline-rich domain (Fig. 4A)(Xu et al., 1995), which can bind SH3-containing proteins (Yu et al., 1994) including Akt (Fig. 4A)(Koral and Erkan, 2012). Inhibition of Dab2 *in vivo* by the electroporation of a dominant negative corresponding to Dab2’s proline-rich domain, markedly reduced Akt association with USP1-TSC1 complex (Fig. 4A), and enhanced Akt phosphorylation during fasting (Fig. 4C). Therefore, Dab2 proline rich domain appears to be essential for Akt inactivation in the absence of insulin or growth factors. Under such physiological and pathological conditions where PI3K-Akt signaling is low (e.g. fasting, type-2 diabetes), liberation of Akt SH3 domain from signaling components that bind and activate Akt may be essential for Akt association with Dab2 and USP1.

Mice lacking USP1 are small, tend to die early, and exhibit multiple defects, including osteopenia (Williams et al., 2011), infertility, and genome instability (Kim et al., 2009). These observations differ from the present findings on the effects of selective downregulation of USP1 in adult wild type mouse muscle. Presumably, the complete deficiency of USP1 during development causes multiple systemic defects that are probably related to dysregulation of DNA repair mechanisms. Because USP1 is an important regulator of PI3K-Akt signaling in adult wild type mice muscles, mice lacking USP1 throughout development seem an inappropriate system to explore the physiological adaptation to prolonged starvation and regulation of PI3K-Akt signaling under normal conditions. Here, we addressed such questions using transient electroporation to reduce USP1 levels or function in normal adult muscles to avoid possible developmental effects.

The expression of USP1 rises in several types of cancer (Williams et al., 2011), where this enzyme probably enhances cell survival by regulating DNA repair processes. However, mutations in USP1 have also been reported in certain human cancers (García-Santisteban et al., 2013), although the functional consequences of these mutations remain elusive. Perhaps these pathological sequelae may be due in part to altered signaling through the PI3K–Akt– FoxO pathway. If these mutations in USP1 cause a gain or loss of enzymatic function, then they might cause dysregulation of PI3K-Akt signaling, which represents a common event in many cancers. The present findings would predict that a deficiency of USP1 could lead to excessive tissue growth and possibly inappropriate activation of PI3K–Akt–FoxO signaling. However, the cellular alterations in these cancers and the effects of the different mutations on USP1 activity are unclear (García-Santisteban et al., 2013), and require in-depth study based upon the present identification of Akt as a new substrate and physiological roles. Because USP1 is a critical regulator of PI3K-Akt signaling, it should also affect cell polarity, cell division, cell adhesion, mobility, metastasis, and neuronal function, and therefore is likely to have important physiological or pathological effects in addition to the ones described herein.

## MATERIALS AND METHODS

### In vivo transfection

All animal experiments were consistent with the ethical guidelines of the Israel Council on Animal Experiments, and the Technion Inspection Committee on the Constitution of the Animal Experimentation. Specialized personnel provided animal care in the Institutional Animal facility. *In vivo* electroporation experiments were performed in adult CD-1 male mice (∼30g)(Envigo) as described previously (Aweida et al., 2018). In brief, 20 µg of DNA plasmid was injected into adult mouse TA muscles, and a mild electric pulse was applied using two electrodes, which were place underneath and on top of the muscle (12V, 5 pulses, 200-ms intervals). The plasmids also encoded GFP to detect transfected muscle fibers. For *in vivo* CRISPR experiment (Fig. 4G), Cas9 encoding plasmid and three guide RNAs against USP1 (Table S2) were co-electroporated into mouse muscle. In fasting experiments food was removed from cages 5 d after electroporation.

### Fiber size analysis

For fiber size analysis, cross-sections of transfected muscles were fixed in 4% PFA, and the fiber membrane stained with laminin antibody. Cross-sectional area of at least 500 transfected fibers (express GFP) and 500 adjacent non-transfected ones in the same muscle section was measured using Imaris 8.2 software (Bitplane), and data collected from at least 4 mice were plotted on a graph. Individual fiber size was determined in the entire muscle cross section by laminin staining (using a 1:50 dilution of laminin antibody and a 1:1,000 dilution of Alexa Fluor 568–conjugated secondary antibody). Images were collected using a Nikon Ni-U upright fluorescence microscope with Plan Fluor 20× 0.5-NA objective lens and a Hamamatsu C8484-03 cooled CCD camera, at room temperature.

### Plasmid DNA and generation of shRNA

The shRNA oligo against USP1 (Table S2) and Lacz were designed using Invitrogen’s RNAi designer tool and were cloned into pcDNA 6.2- GW/EmGFP-miR vector using Invitrogen’s BLOCK-iT RNAi expression vector kit. The GFP-tagged USP1^(C90S)^ dominant-negative encoding construct has been described previously (Olazabal-Herrero et al., 2016). The mammalian expression vector encoding the C-terminal proline rich domain of Dab2 was obtained from Dr. Philip Howe (Cleveland Clinic Lerner College of Medicine, Ohio, USA)(Hocevar et al., 2005), and the one encoding SFB-tagged UAF1 were from Dr. Maddika Subbareddy (CDFD, India)(Gangula and Maddika, 2013). The plasmid encoding Akt^(K8R)^ has been described previously (Yang et al., 2009). The Cas9 encoding plasmid was obtained from IDT (1072566), and the three USP1 guide RNAs were purchased from Transomic.

### Antibodies and materials

Anti-actin (#A2066), GAPDH (#G8975), and Laminin (#L9393) were purchased from Sigma-Aldrich. Anti-phospho Insulin receptor (Y1361)(#60946) was from Abcam; anti-USP1 (#14346-1-AP) was from Proteintech; anti-UAF1 (#514473) and anti-Dab2 (#136964) were from Santa cruz; anti-Akt (#9272), FoxO (#2497), Insulin receptor (#5025), phospho Akt T308 (#9275), phospho FoxO (#9464), phospho GSK3 (#8566), phospho S6K (#9205), TSC1 (#6935), and TSC2 (#3990S) were from Cell Signaling Technology. Anti-polyubiquitin K63-linked chains was from Millipore (#051308). The secondary antibodies anti-rabbit IgG-HRP (#111-035-144) and anti-mouse IgG-HRP (#115-035-003) were from Jackson ImmunolResearch, and Alexa Flour 568 goat anti-rabbit IgG was from ThermoFisher (#A-11011). The DUB scan kit was from Ubiquigent (#67-0006-001).

### Skeletal muscle homogenization

Skeletal muscles were homogenized in cold Homogenization Buffer (20 mM Tris pH 7.2, 5 mM EGTA, 100 mM KCl, 1% Triton X-100, 10 mM Sodium Pyrophosphate, 50 mM NaF, 2 mM Sodium Orthovanadate, 10 ug/ml Aprotinin, 10 ug/ml Leupeptin, 3 mM Benzamidine, 1 mM PMSF, 1% Phosphatase Inhibitor Cocktail (Sigma #P0044)). Following incubation for 1 hr at 4°C, and centrifugation at 6000 × g for 20 min at 4°C, the supernatant (soluble fraction) was stored at −80°C.

### Protein analysis

For immunoblotting, soluble fractions from TA muscles, as well as the *in vitro* deubiquitination reactions, were resolved by SDS-PAGE, transferred onto Nitrocellulose or PVDF membranes, and immunoblotted with specific antibodies and secondary antibodies conjugated to HRP. Immunoprecipitation experiments from muscle homogenates (1:50 dilution of primary antibody) were performed overnight at 4°C and then protein A/G agarose beads was added for 3 hr incubation at 4°C. To remove nonspecific or weakly associated proteins, tubes were centrifuged at 2500 × rpm for 5 min at 4°C, supernatant was discarded and precipitates were washed extensively with 10 bed volumes of high (50m M Tris-HCl pH 8, 500 mM NaCl, 0.1% SDS, 0.1% Triton, 5 mM EDTA), medium (50 mM Tris-HCl pH 8, 150 mM NaCl, 0.1% SDS, 0.1% Triton, 5 mM EDTA), and low (50 mM Tris-HCl pH 8, 0.1% Triton, 5 mM EDTA) salt buffers. Protein precipitates were then analyzed by SDS-PAGE and immunoblotting.

For glycerol gradient fractionation, two glycerol stocks of 10% and 40% were prepared in buffer G (20 mM Tris pH 7.6, 0.46 mM EDTA/NaOH pH 8, 100 mM KCl, 1 mM DTT, 0.25% Sodium deoxycholate, 1 mM Sodium Orthovanadate), layered one on top of the other, and the gradient was left to form overnight at 4°C. Muscle homogenates from fed and fasted mice (300ug) were loaded on the top of the gradient, and following centrifugation for 24 hr at 35,000 rpm (SW 55Ti rotor) and 4°C, equal volumes of protein fractions were collected from the bottom of the tube. Proteins were precipitated using 10% Trichloroacetic Acid at 4°C overnight, centrifuged at 14,000 rpm for 10 min at 4°C, washed twice with acetone, and alternate fractions analyzed by SDS-PAGE and immunoblotting.

### Mass Spectrometry analysis

To identify the proteins mediating USP1-Akt association during fasting, four immunoprecipitation experiments from mouse muscle homogenates during fasting were subjected to in-solution tryptic digest followed by mass spectrometry analysis. An IgG antibody was used as a control for non-specific binding. The data was processed using the Genedata Expressionist system and searched using Mascot against the mouse sequences in UniprotKB appended with 125 common lab proteins (like keratins, albumin ect.). Data was filtered for 1% FDR at the peptide and protein level. The ratios of protein intensities in each sample vs. IgG control were calculated, and proteins that were at least 3 times more abundant in the immunoprecipitation reaction than in IgG control were considered. Among identified proteins, 29 proteins were enriched in all four samples and were at least 1000 times more abundant in the immunoprecipitation samples than in IgG control.

### Glucose tolerance test

D-glucose (1 mg/gr body weight) was administered to mice by intraperitoneally injection. Blood glucose levels were measured before the injection, and 5, 15, 30, 45, 60, 75, and 120 min later by blood glucose monitor device.

### In vivo measurement of protein synthesis rates

Five days after muscle electroporation, mice were deprived of food for 2 d, and 30 min before euthanasia, they were anesthetized and injected with puromycin (0.04 µmol/g body weight, i.p.). Exactly 25 min after puromycin injection, mice were sacrificed, and dissected TA muscles were snap-frozen in liquid nitrogen. Effects on protein synthesis rates were assessed by SDS-PAGE and immunoblotting using puromycin antibody (Millipore, #MABE343).

### Quantitative real-time PCR

Total RNA was isolated from muscle using TRI-reagent (Sigma, #T9424) and served as a template for synthesis of cDNA by reverse transcription (Quanta script cDNA synthesis kit, #84005, #84002). Real-time qPCR was performed on mouse target genes using specific primers (Table S2) and PerfecTa SYBR Green FastMix (Quanta 84071) according to manufacturer’s protocol.

### Statistical analysis and image acquisition

Data are presented as means ± SEM. The statistical significance was determined with one-tailed paired Student’s t test. Alpha level was set to 0.05. Black and white images were processed with Adobe Photoshop CS5, version 12.1. Quantity One algorithm (Bio-Rad Laboratories version 29.0) was used for densitometric measurements of protein bands intensity.

## ACKNOWLEDGMENTS

This project was supported by the Israel Science Foundation (grant no. 623/15), the Israel Ministry of Science, Technology and Space (grant no. 2015-3-74), and the Niedersachsen-Deutsche (grant no. ZN3008) grants to S. Cohen. Additional funds have been received from the Russell Berrie Nanotechnology Institute, Technion, and Dr. Bernard and Bobbie Lublin Cancer Research award to S. Cohen.

We acknowledge the de Botton Institute for Protein Profiling at Weizmann Institute of Science for the mass spectrometry analysis.

## ABBREVIATIONS LIST

USP1: Ubiquitin-Specific Protease 1
Dab2: Disabled-2
DUB: Deubiquitinating enzyme
TA: Tibialis Anterior
DN: Dominant Negative

## SUPPLEMENTAL INFORMATION TITLES AND LEGENDS

**Figure S1. USP1 and UAF1 are not induced during fasting.**

Quantitative RT-PCR of mRNA preparations from muscles from fed and fasted mice using primers for USP1 and UAF1. Data are plotted as the mean fold change relative to fed control. n = 6.

**Figure S2. During fasting, downregulation of USP1 stabilizes TSC1 and increases levels of phosphorylated Akt.**

(A) Inhibition of USP1 does not affect TSC1 expression during fasting. Quantitative RT-PCR of mRNA preparations from atrophying and control muscles expressing shLacz or USP1^(C90S)^ using primers for TSC1. Data are plotted as the mean fold change relative to fed control. n = 4. *, P < 0.05 vs. shLacz in fed.

(B-C) Soluble fraction of TA muscles expressing shLacz or shUSP1 from fed and fasted (for 24hr, 36hr or 48hr) mice were analyzed by SDS-PAGE and immunoblotting.

**Figure S3. Dab2 is not induced during fasting.**

Quantitative RT-PCR of mRNA preparations from muscles from fed and fasted mice using primers for Dab2. Data are plotted as the mean fold change relative to fed control. n = 4.

**Table S1.**
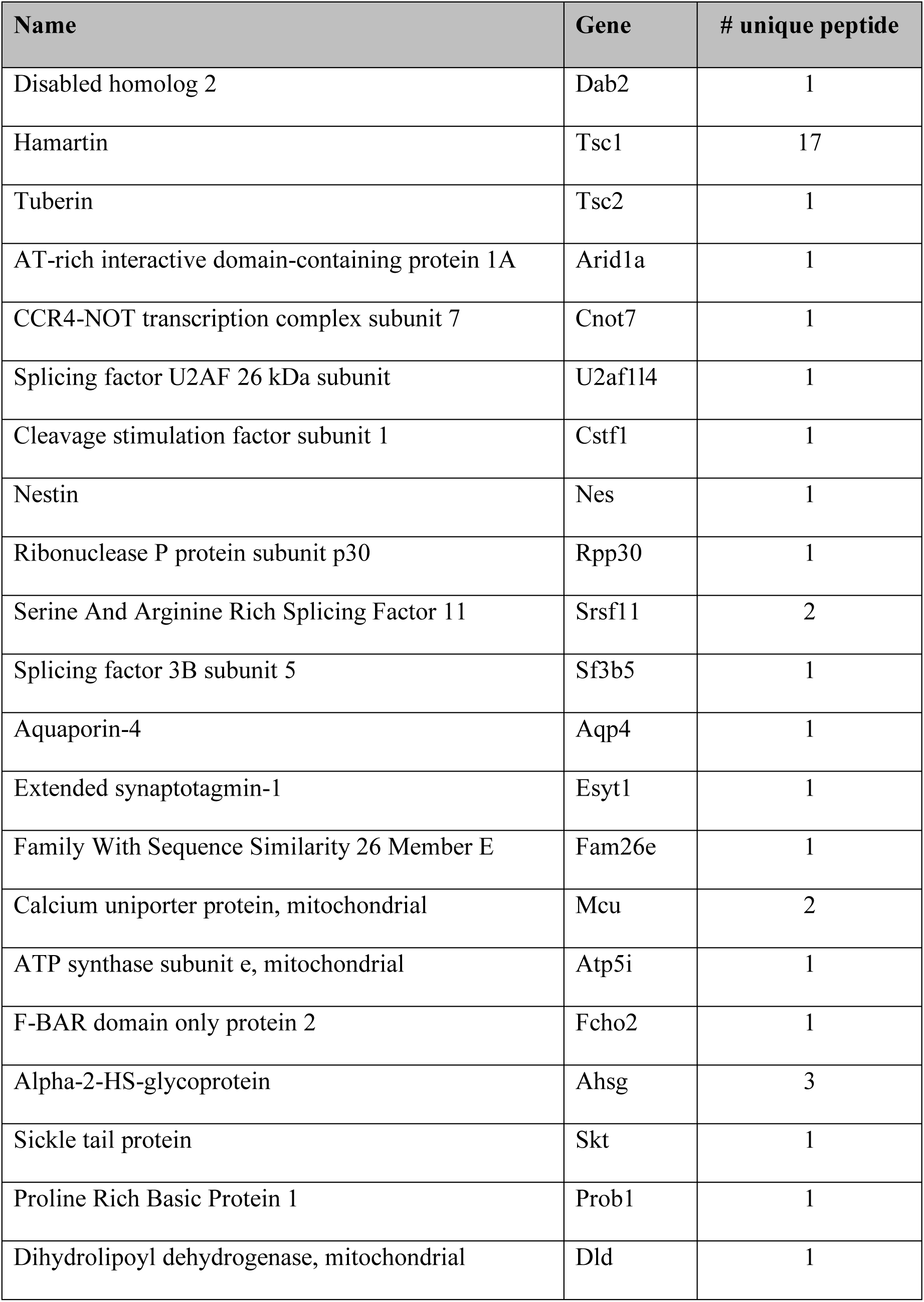
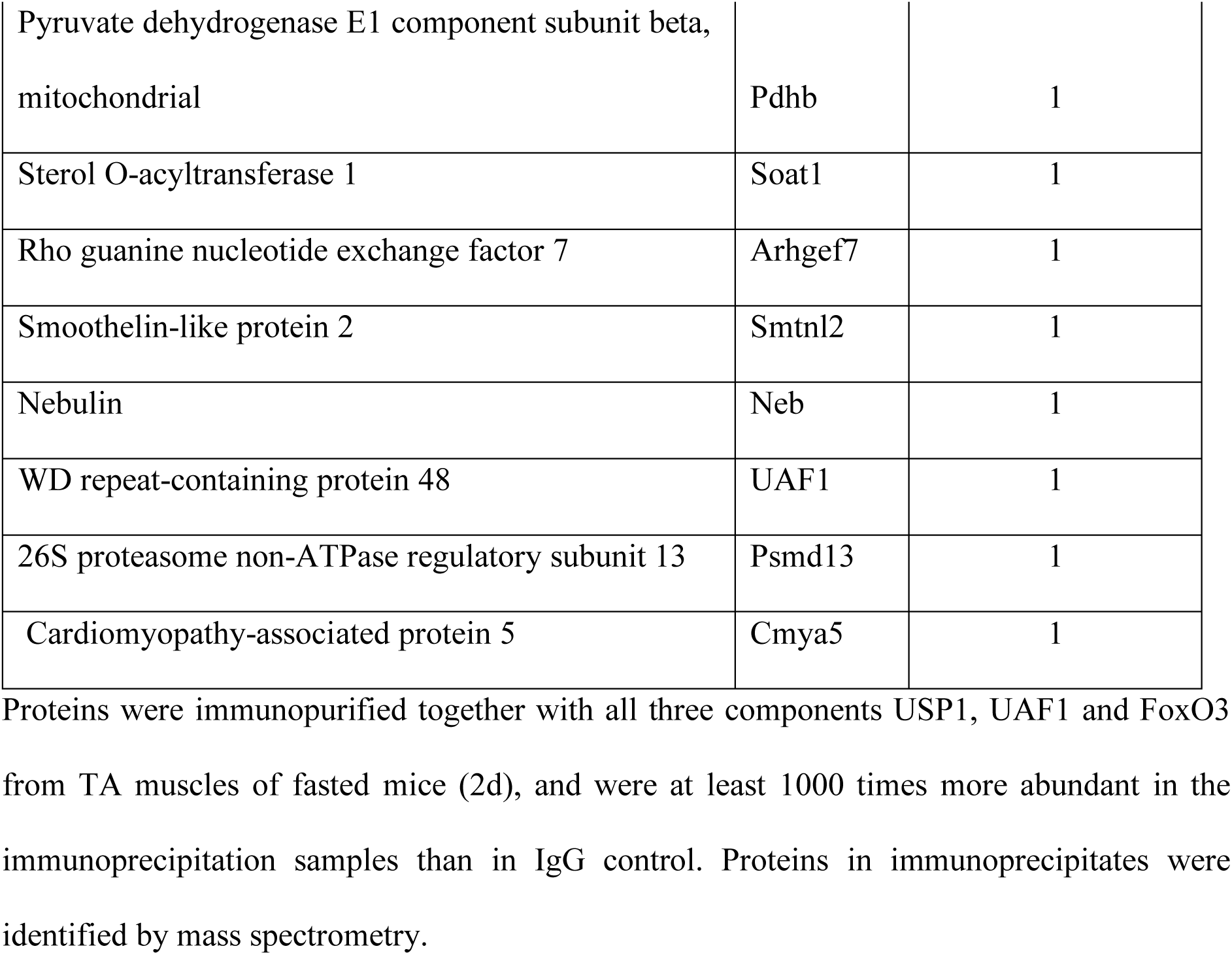
Proteins that bound USP1/UAF1 and FoxO3 in muscle homogenates from fasted mice (2d).

**Table S2.**
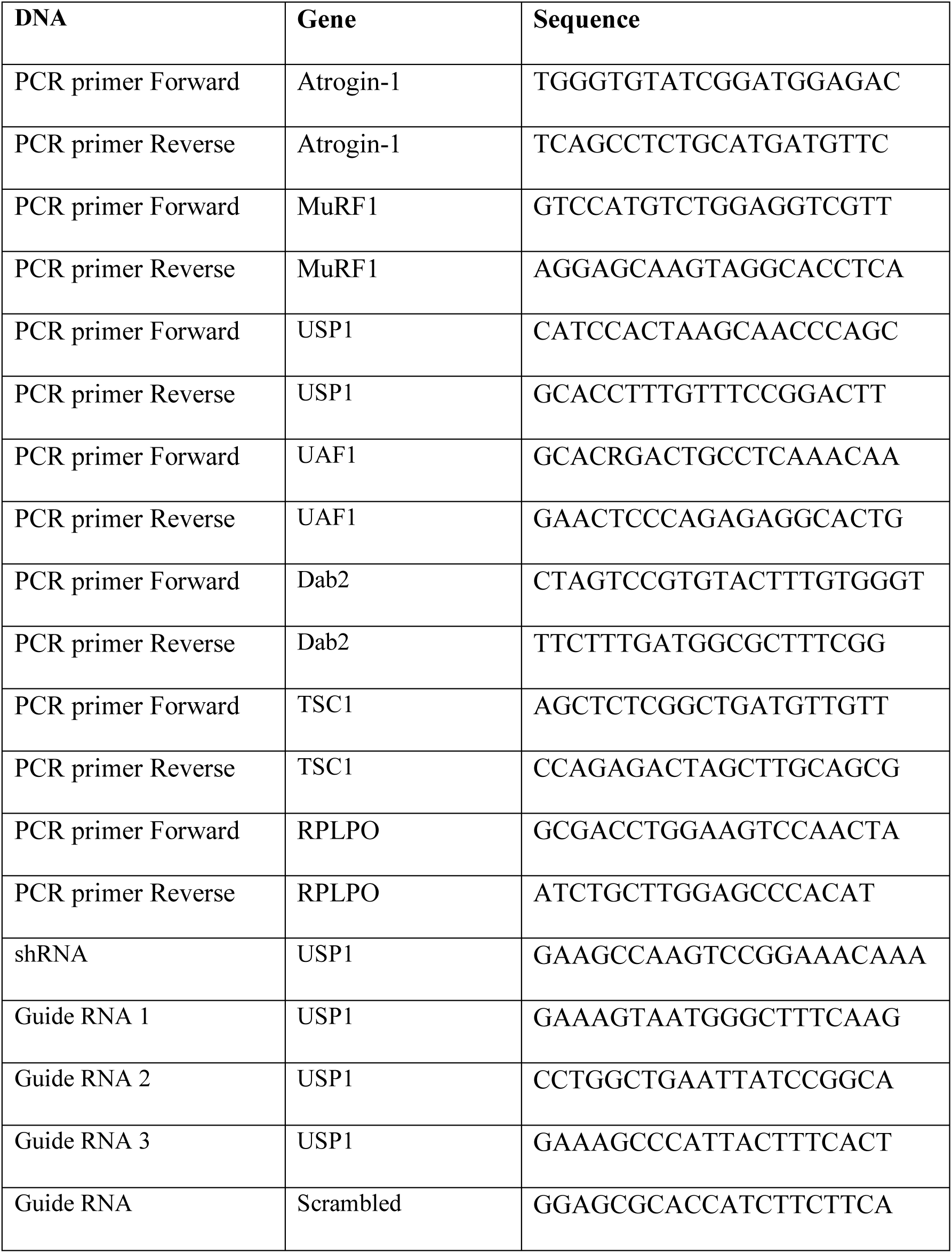
qPCR primers, shRNA oligos, and guide RNAs used in the present study.

