## Supplementary figures and images for "USP1 deubiquitinates protein kinase Akt to inhibit PI3K-Akt-FoxO signaling"

### Supplemental figures

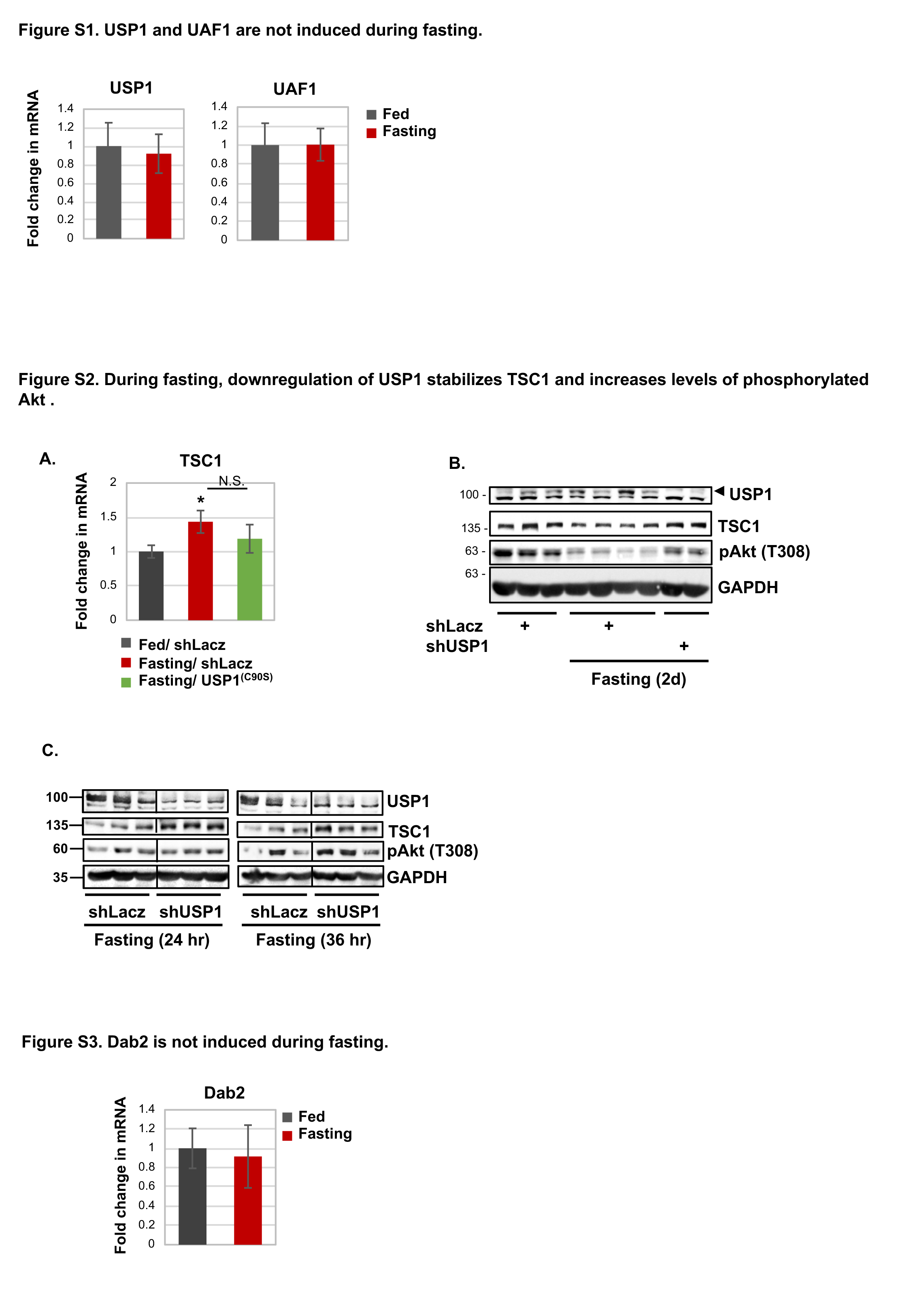
